# A curated collection of human vaccination response signatures

**DOI:** 10.1101/2021.04.15.439017

**Authors:** Kenneth C. Smith, Daniel G. Chawla, Bhavjinder K. Dhillon, Zhou Ji, Randi Vita, Eva C. van der Leest, Jing Yi (Jessica) Weng, Ernest Tang, Amani Abid, The Human Immunology Project Consortium (HIPC), Bjoern Peters, Robert E.W. Hancock, Aris Floratos, Steven H. Kleinstein

## Abstract

Recent advances in high-throughput experiments and systems biology approaches have resulted in hundreds of publications identifying “immune signatures”. Unfortunately, these are often described within text, figures, or tables in a format not amenable to computational processing, thus severely hampering our ability to fully exploit this information. Here we present a data model to represent immune signatures, along with the Human Immunology Project Consortium (HIPC) Dashboard (www.hipc-dashboard.org), a web-enabled application to facilitate signature access and querying. The data model captures the biological response components (e.g., genes, proteins, cell types or metabolites) and metadata describing the context under which the signature was identified using standardized terms from established resources (e.g., HGNC, Protein Ontology, Cell Ontology). We have manually curated a collection of >600 immune signatures from >60 published studies profiling human vaccination responses for the current release. The system will aid in building a broader understanding of the human immune response to stimuli by enabling researchers to easily access and interrogate published immune signatures.

## Introduction

Systems-level profiling of the human immune system has generated important insights into the mechanisms by which humans respond to exposures such as vaccination. These studies, including many conducted through the Human Immune Project Consortium (HIPC), have generated hundreds of publications. While repositories exist to promote re-use of primary experimental immunology data generated from these efforts, such as the Gene Expression Omnibus^1^ (GEO) and the NIAID Division of Allergy, Immunology, and Transplantation (DAIT)-sponsored Immunology Database and Analysis Portal^2^ (ImmPort), there is no centralized framework to aggregate and organize the published findings resulting from the analysis of this data, and particularly the coherent sets of biomarkers, termed here “signatures”. Additionally, such signatures are not published in a consistent format between publications and may be presented as text, tables, or images. This heterogeneity presents a barrier to comparative analyses since identifying published signatures, for example of a vaccine response, requires extensive manual curation of the literature that must be repeated by investigators each time they wish to interpret a set of results. Here, we propose a model to standardize the representation of these published findings and present the Human Immunology Project Consortium (HIPC) Dashboard—a searchable interface to query curated signatures from the corpus of human immunology literature.

We define a ‘signature’ as the information required to specify a published result. This includes alterations in the levels of a set of one or more response component(s), i.e., biological entities such as genes or cell types, that are defined by a particular comparison in the context of an immune exposure. The signature also includes contextual information (termed metadata) such as the conditions and circumstances under which the signature was identified, the tissues or cells that were assayed, as well as clinical data such as demographic information about the groups that were included in the analysis. As a motivating example, a study by Bucasas et al.^3^ identified a set of genes that are up-regulated in individuals with higher antibody responses (comparison) after vaccination with the 2008-2009 trivalent influenza vaccine (exposure) in an adult cohort. The expression of genes STAT1, IRF9, SPI1, CD74, HLA-E, and TNFSF13B one day after influenza vaccination was predictive of greater antibody responses. In the paper, these results were represented in a table, though similar findings often appear as text or within figures. Without standardization, such findings are not easily accessible to the wider scientific community for further analysis.

Several existing resources define pathway and gene module signatures through reanalysis of raw data, but few capture the original findings published with these data or are specifically geared towards human immunology research. Among these resources are the OMics Compendia Commons (OMiCC)^4^, EnrichR^5,6^, the integrative Library of Integrated Network-based Cellular Signatures (iLINCS)^7^, the Molecular Signatures Database (MSigDB)^8,9^, and VaximmutorDB^10^. OMiCC crowdsources annotations for gene expression data to be used in reanalysis and novel signature generation. EnrichR and iLINCS offer biological annotations built from data reanalyzed en masse, but similarly do not capture published findings. MSigDB does include manually curated gene signatures along with those derived from data reanalysis, albeit with fewer contextual details than captured for the HIPC Dashboard. VaximmutorDB captures published gene expression and proteomic signatures but not cell-type frequency signatures, and signatures from this database are not yet downloadable in a machine-readable format.

To improve access and to promote reuse of published signatures, we designed a data model that standardizes the content and context of published immune signatures. Our initial curation efforts have focused on gene expression and cell-type frequency/activation signatures of human vaccine responses, but this framework is extensible to other domains such as response to infection. We captured what is changing, (e.g. groups of genes), how that response component changed (e.g. up- or down-regulation), where this change was observed (e.g. in sorted CD8+ T cells from adults), and the comparison that was performed (e.g. individuals with high vs. low antibody titers post-vaccination). We then manually curated signatures from publications both within and outside of HIPC that described changes in gene expression, cell-type frequencies, or cell activation state in response to vaccination. To disseminate these immune signatures, we developed the HIPC Dashboard (www.hipc-dashboard.org), a web-accessible, user-friendly interface to enable signature searching and browsing, and to facilitate rapid comparative analyses. The design of the HIPC Dashboard is based on a similar infrastructure we developed previously for the Cancer Target Discovery and Development network, the CTD^2^ Dashboard^11^, and leverages the same underlying ontological framework for the standardized representation of research findings as well as the emphasis on the consistent, curation-mediated use of controlled vocabularies for linking findings reported in different publications.

## Results

### A Data Model for Immune Signatures of Vaccination

We developed a data model that captures, in a detailed and consistent format, the essential information embedded in published immune signatures of vaccination for dissemination through the HIPC Dashboard (**Table 1)**. Key elements of this data model (e.g., genes, vaccines, etc.) are specified using controlled vocabularies, thus making immune signatures of vaccination amenable to data mining and promoting compatibility with projects both within and outside of HIPC. A signature as defined in this model encapsulates both a change in the behavior or abundance of a biological response component as well as the metadata describing the context under which the signature is identified, including (1) the tissue in which the signature was observed, (2) the immune exposure and timing underlying the observed comparison, and (3) clinical details of the cohort from which tissue samples were taken, including age (**Figure 1**). The model accommodates many types of biological response components (gene, protein, metabolite, pathway, and cell type (e.g. subsets of blood cells). We focused on gene expression and cell type signatures of vaccination, but the data model and HIPC Dashboard infrastructure are flexible and can be easily expanded to accommodate arbitrary signature types.

**Table 1.** Data model for capturing immune signatures. Genes and cell types are captured as response components, with terms standardized against NCBI/HGNC or CL+PRO, respectively. Exposures are captured and standardized against VO terms and NCBI Taxonomy IDs. Metadata includes observed tissue, study timing, cohort descriptors, and age characteristics. *fields in the Immune Exposure model^17^

| Elements | Content Definition | Field value Example | Ontology | Data Type | Content Format |
| --- | --- | --- | --- | --- | --- |
| **Response Component** | The biological entity being observed: either a specific gene or cell subset | CKS1B | NCBI/HGNC for genes, Cell Ontology for cell types, PRO for proteins) | String | Controlled |
| **Response Component Type** | Type of response agent, e.g. gene, cell subset | gene |  | String | Fixed List |
| **Tissue Type** | The type of cells analyzed, e.g. whole blood, gated cells | whole blood (UBERON:0000178) | Blood + PRO/CL | String | Controlled |
| **Exposure Process Type \*** | Category of immune exposure | Vaccination |  | String | Fixed List |
| **Exposure Material \*** | Eg, the type of vaccine administered | VO:0001176 | VO | String | Controlled |
| **Disease Name \*** | The condition being observed | None | DO | String | Controlled |
| **Disease Stage \*** | Stage of infection (e.g. acute, chronic, post, etc.) | None | OGMS | String | Controlled |
| **Target Pathogen** | The pathogenic organism (e.g. virus, bacterium, prion, fungus) being studied | Influenza A, Influenza B | NCBI Taxonomy ID | String | Controlled |
| **Vaccine Year** | Year for seasonal vaccines | 2012 |  | Numeric | Free text |
| **Adjuvant** | A substance added to vaccines to increase the body's immune response to the vaccine |  | VO | String | Controlled |
| **Route** | Eg oral, nasal spray, intradermal injection, etc. | Intramuscular, intradermal, transcutaneous | VO | String | Controlled |
| **Scheduling** | The number of times a substance is administered within a specific time period. | e.g. prime / boost scheme |  | String | Free text |
| **Time Point** |  | 1 |  | Numeric |  |
| **Time Point Units** |  | day |  | String | Fixed List |
| **Baseline Time Event** | The starting point against which other events are compared | time of vaccination (first dose) |  | String | Free Text |
| **Cohort** | Features that describe study cohorts | e.g. antibody responders |  | String | Free Text |
| **Age Min** |  | 18 |  | Numeric |  |
| **Age Max** |  | 45 |  | Numeric |  |
| **Age Units** | {days, weeks, months, years} | years |  | String | Fixed List |
| **Publication Reference** | PubMed unique identifier of an article. | 30843873 |  | Numeric | Controlled |
| **Publication Year** | The year in which the study was published | 2019 |  | Numeric |  |
| **Publication Reference URL** | Link to article in PubMed | https://www.ncbi.nlm.nih.gov/pubmed/30843873 |  | String | URL |
| **Comparison** | The contrast used for deriving the signature | 1d-0d |  | String | Free Text |
| **Response Behavior** | Observed change in the response agent under the comparison | Up |  | String | Free Text |

**Figure 1.**
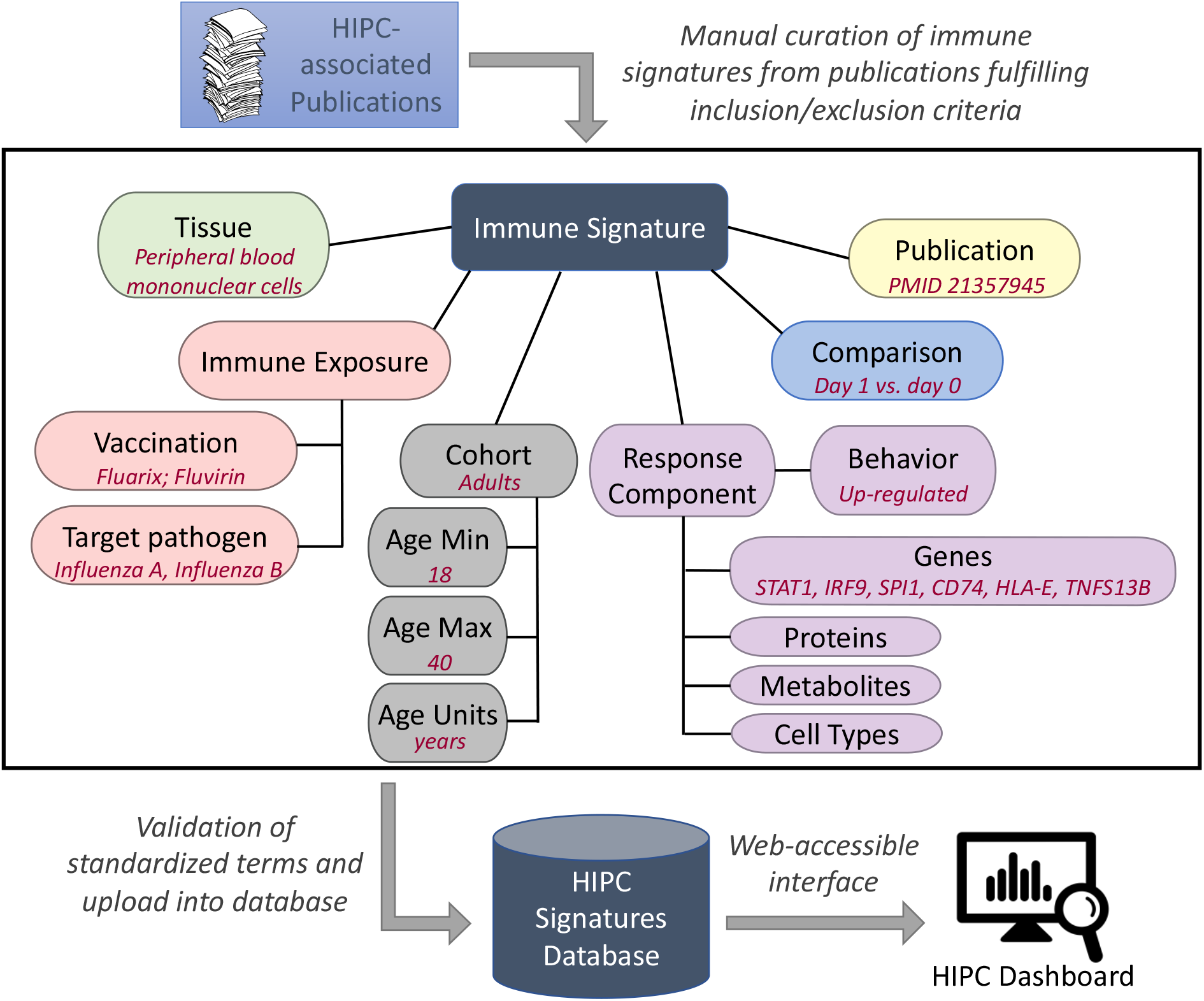
Overview of the manual curation process for extracting immune signatures from relevant publications into the HIPC signatures database and a web-accessible HIPC Dashboard. The middle panel highlights the various fields that are captured for a given immune signature, with examples provided in red font. Key fields are standardized using existing ontologies or pre-defined criteria in order to capture a wide array of signatures.

To facilitate data mining and comparative analyses between different conditions including vaccine types, and to afford consistency between this database and other projects using the same controlled vocabulary terms, standardized terms and ontology links were used for as many biological response components, immune exposures, and demographic fields as possible. Gene and cell type response components were standardized to the HGNC^12^ (as provided through the NCBI) and Cell Ontology^13,14^ (CL) respectively to enable comparisons across publications using different naming conventions. Cell types from the Cell Ontology were further differentiated using protein marker terms drawn from Protein Ontology^15^ (PRO), where possible (e.g. IFNG+ T cells). This same naming convention is used to describe the tissue in which the signature was observed^16^.

To annotate the immune challenges driving each signature, we utilized the Immune Exposure model^17^, which provides a standardized description of a broad range of potential and actual exposures to different immunological agents (e.g., vaccination, laboratory confirmed infection, living in an endemic area, etc.). Immune exposures are broken down into Exposure Process, Exposure Material, Disease Name, and Disease Stage. Each of these components is modeled using standardized ontology terminology. Within the data model for the HIPC Dashboard, Exposure Materials such as vaccines are captured using terms in the Vaccine Ontology^18^ (VO), which further link to target pathogens and strains using the NCBI Taxonomy^19,20^. While these ontology choices reflect our initial focus on vaccination, the data model can accommodate other exposure processes beyond vaccination, with links to appropriate ontologies. Integration of the Immune Exposure model in the HIPC Dashboard data model promotes interoperability with other projects that have adopted its use, both within and outside of HIPC, including data repositories such as ImmPort and the Immune Epitope Database^21^.

Cohort information that is important for interpreting signatures is also captured. Cohort descriptors can vary widely between studies and can include, for example, sex, antibody response titers, geographic location, health status, vaccination or infection history, etc. This information is currently recorded as unstructured text to maintain flexibility. Cohort age range is standardized separately by storing minimum and maximum ages along with their units. Additional fields describe the particular perturbations that drive the changes to the biological response components. The “comparison” field describes the cohort groups whose differential response under the perturbation is measured. Examples of comparison groups include measurements taken at two different time points (e.g. day1 vs. day 0), correlation with antibody response, differing antibody response outcomes (e.g. high vs. low responders), or comparisons across different demographic parameters such as age or sex (e.g. younger vs. older, female vs. male). The “response_behavior” field captures the directionality of the differential response (e.g. up or down, positively- or negatively-correlated) under the specified comparison. Fields for which a formal controlled vocabulary was not used, such as cohort descriptions and the comparison, are stored as free text.

Finally, each signature is tagged with a Pubmed ID (PMID) and publication year field, to connect observations to their literature sources.

### Manual Curation of Published Signatures

A defined Pubmed search strategy was used (see Methods) to assemble an initial list of publications comprising studies that involved a systems-level profiling of measurable changes before and after vaccination in human subjects. The publications were culled for signatures reporting statistically significant changes in gene expression, cell-type frequency or cell activation state induced by an immune exposure when comparing groups with different features (such as high vs low responders as defined by antibody titers). We focused on components that also included information about response behavior (e.g. up- or down-regulated). In total, 665 immune signatures were manually curated from 69 published studies. After standardization and quality control (see Methods), these curated gene and cell type signatures included 13,812 unique genes, 152 unique cell types (including protein markers and additional type-modifiers), and 44 pathogens across 56 vaccines **(Table 2)**. Table 3 illustrates a typical gene-expression type signature after tissue, gene symbol and pathogen standardization.

**Table 2.** Dashboard summary statistics for gene and cell-type signatures. “Joint” refers to the union of the two signature types, as they overlap in the various components. “Total Response Components” lists the number of genes or parent cell types from the Cell Ontology (CL) across all signatures. When additional cell-type markers are included, e.g. from the Protein Ontology (PRO), there are 152 unique cell types represented among the signatures. “Response Components per Signature” shows the range of the number of response components found in individual signatures.

| Signature Type | Vaccines | Target Pathogens | Publications | Signatures | Total Response Components | Response Components per Signature |
| --- | --- | --- | --- | --- | --- | --- |
| Gene | 52 | 38 | 62 | 480 | 13,812 genes | 1 to 2,036 genes |
| Cell type | 28 | 26 | 31 | 185 | 47 cell types | 1 to 9 cell types |
| Joint | 56 | 44 | 69 | 665 |  |  |

**Table 3.** Key fields in the immune signature data model for a gene expression signature. This signature reports positive correlation between gene expression and a computed titer response index (TRI).

| Column name | Ontology | Values |
| --- | --- | --- |
| response_component | HGNC | STAT1, IRF9, SPI1, CD74, HLA-E, TNFSF13B |
| tissue_type | Cell Ontology | peripheral blood mononuclear cell |
| exposure_material | Vaccine Ontology | VO:0000045; VO:0000046 (Fluarix; Fluvirin) |
| target_pathogen | NCBI Taxonomy | Influenza A virus (A/Brisbane/59/2007(H1N1));<br>Influenza A virus (A/Brisbane/10/2007(H3N2));<br>Influenza B virus (B/Florida/4/2006) |
| vaccine_year |  | 2008 |
| time_point |  | 1 |
| time_point_units |  | days |
| baseline_time_event |  | time of vaccination (first dose) |
| cohort |  | 18-40 yo, subgroup high responders |
| publication_reference (PMID) |  | 21357945 |
| comparison |  | correlated with titer response index (TRI) |
| response_behavior |  | positive |

### The HIPC Signatures Dashboard

The data model provides a means for representing immune signatures in a structured, standardized, machine-readable manner while the curation process enables the cross-referencing of signatures from different publications based on their shared response components, by enforcing the consistent use of controlled vocabularies for codifying these components (e.g., genes, cell types, tissues, vaccines, and pathogen strains). These capabilities come together in the “HIPC Dashboard” (http://hipc-dashboard.org), a web application developed to enable dissemination of the curated set of immune response signatures. The HIPC Dashboard allows signature browsing, as well as searching for one or multiple response components (using the corresponding controlled vocabulary terms and their synonyms), to retrieve all immune signatures involving the query response component(s) across all curated publications.

The central viewable element of the HIPC Dashboard is the “Observation Summary”, a human-readable description of the information captured in an immune signature. Observation summaries are constructed “on the fly”, using template text devised as part of the curation process. The template has placeholders for the various elements of an immune signature, including the response component (gene or cell type) and the response behavior type (up/down or correlation). When a specific signature is selected in the process of browsing or searching the Dashboard, the observation summary for that signature is instantiated by replacing the template placeholders with the relevant values from that signature. For example, a joint search on the terms “CD4” and “Zostavax” yields about 35 observation summaries. One of these is related to a change in cell-type frequency:

In peripheral blood mononuclear cell, CD4-positive, alpha-beta memory T cell **& CD38+, HLA-DR+, VZV tetramer+** frequency was **up** at **14 days** from **time of vaccination** for the comparison **14d vs 0d** in cohort **50-75 yo** after exposure to Zostavax targeting Human alphaherpesvirus 3 (**details_»**)

Here, the “&” sign separates the Cell Ontology cell type from Protein Ontology surface and other markers. In a second example, a joint search on the terms “CXCL10” and “BCG” yields 6 observation summaries, one of which reports a correlation of gene expression (at 1 day post-vaccination) to an ELISpot result at 28 days post-vaccination:

In blood, CXCL10 gene expression at **1 day** from **time of vaccination** was **positively correlated** with **IFN-gamma ELISpot spot forming cell 28d** in cohort **4-6 mo subgroup BCG-primed** after exposure to MVA85A targeting Mycobacterium tuberculosis variant bovis BCG (**details »**)

In both cases, placeholders in the observation summary template have been replaced by controlled terms for the response components and ontology-linked metadata (blue, hyperlinked text) and by free-text metadata describing informative experiment details (black, bold text). Following the hyperlink for a controlled term leads to a dedicated page for the corresponding biological entity, providing additional details (including links to relevant external annotation sources, e.g., Entrez, GeneCards and UniProt for genes) as well as a listing of all the immune signatures stored in the Dashboard that involve that entity **(Figure 2A)**. Further, the “details” link at the end of each observation summary points to an “Observation” page **(Figure 2B)** containing detailed information about the corresponding immune signature, including a full listing of all its available metadata. This includes, for example, structured text for values such as age group and days post-immunization, and links to download the full signature source data (including all metadata) in tab-delimited form. Additionally, each observation includes a link to a file containing the complete set of response components from which it was derived, e.g. the full list of genes or cell types.

**Figure 2.**
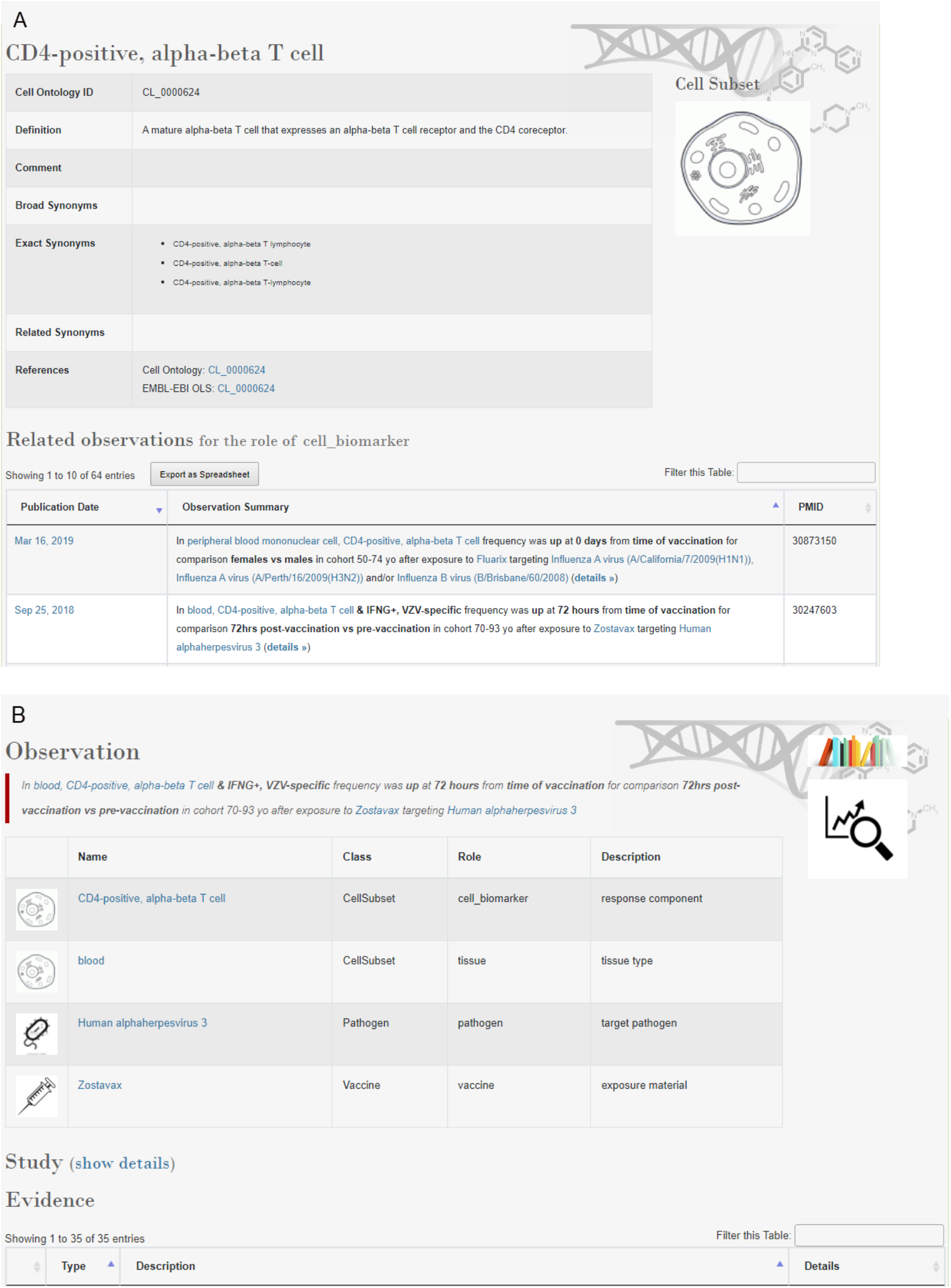
HIPC Dashboard web interface. **A**. Subject page for cell type “CD4-positive, alpha-beta T cell” showing a link-out to the Cell Ontology, the filtering box to further narrow the displayed observations, and the first two observation summaries (“Related observations”). **B**. Partial view of a details page for a CD4-positive, alpha-beta T cell observation. For each controlled term, its name, plus its class, role, and description are shown. Linked pages list details from the relevant ontology and list all observations containing that term. The class equates to its controlled vocabulary type; values are cell subset, gene, pathogen, and vaccine. Roles are used to further differentiate how each term, whether controlled or standardized, is being used. Among the classes in the HIPC Dashboard, only the class “cell subset” has more than one role, these being “tissue” and “cell_biomarker”. Full metadata, not shown here, is contained in the table labeled “Evidence” at the bottom.

## Discussion

Users of the HIPC Dashboard can easily search and examine hundreds of immune signatures related to human vaccination responses. Consolidating these publications, standardizing their findings in a database, and disseminating them through the Dashboard interface allows for rapid comparative analyses and re-use of published findings. This is particularly important for identifying commonalities across studies that may reflect shared mechanisms. The HIPC Dashboard can offer broad insights into the mechanisms by which our immune systems respond to vaccination and will be of great value to the vaccine research community. Although the HIPC Dashboard is not designed as an analysis engine, all signatures are made available for download so that users may perform more sophisticated and targeted downstream analyses.

Among data resources dedicated to the collection of vaccination signatures, the HIPC Dashboard is nearly unique in its emphasis on manual curation of published literature. To the best of our knowledge, only MSigDB and VaximmutorDB maintain signatures curated from publications. MSigDB provides minimally redundant gene sets for enrichment analyses, but unlike the HIPC Dashboard does not attempt to capture the full biological context of published results. A reduced set of our curations has recently been made available for gene set enrichment analyses through MSigDB under the C7 VAX gene sets. VaximmutorDB provides access to a collection of immune factors (genes/proteins) that change in response to vaccination against 46 pathogens. Compared to VaximmutorDB, the HIPC Dashboard offers several advantages, including: (i) a wider breadth in the types of response components and immune changes that are captured, (ii) improved browsing functionality that facilitates comparisons of immune changes across studies and vaccines, and (iii) the ability to download signature data.

As of the date of this publication, 152 unique cell types and 13,812 distinct genes have been collected in the HIPC Dashboard; this large number of published results allows users to quickly examine the role of particular biological response components, such as individual genes or cell types, across studies. Most of the currently curated gene signatures have fewer than 50 genes, with a range of 1 to 2,036 (**Figure 3A**), while most cell-type frequency or cell activation state signatures have only a single cell type, with a range up to 9 (**Figure 3B**). These signatures represent findings from 16 tissues and tissue extracts, including blood, PBMCs, T cells, B cells, monocytes, and NK cells. Nearly 250 entries describe changes over time, more than 75 capture antibody response-associated signatures, and several others come from studies that report effects of age and T cell responses.

**Figure 3.**
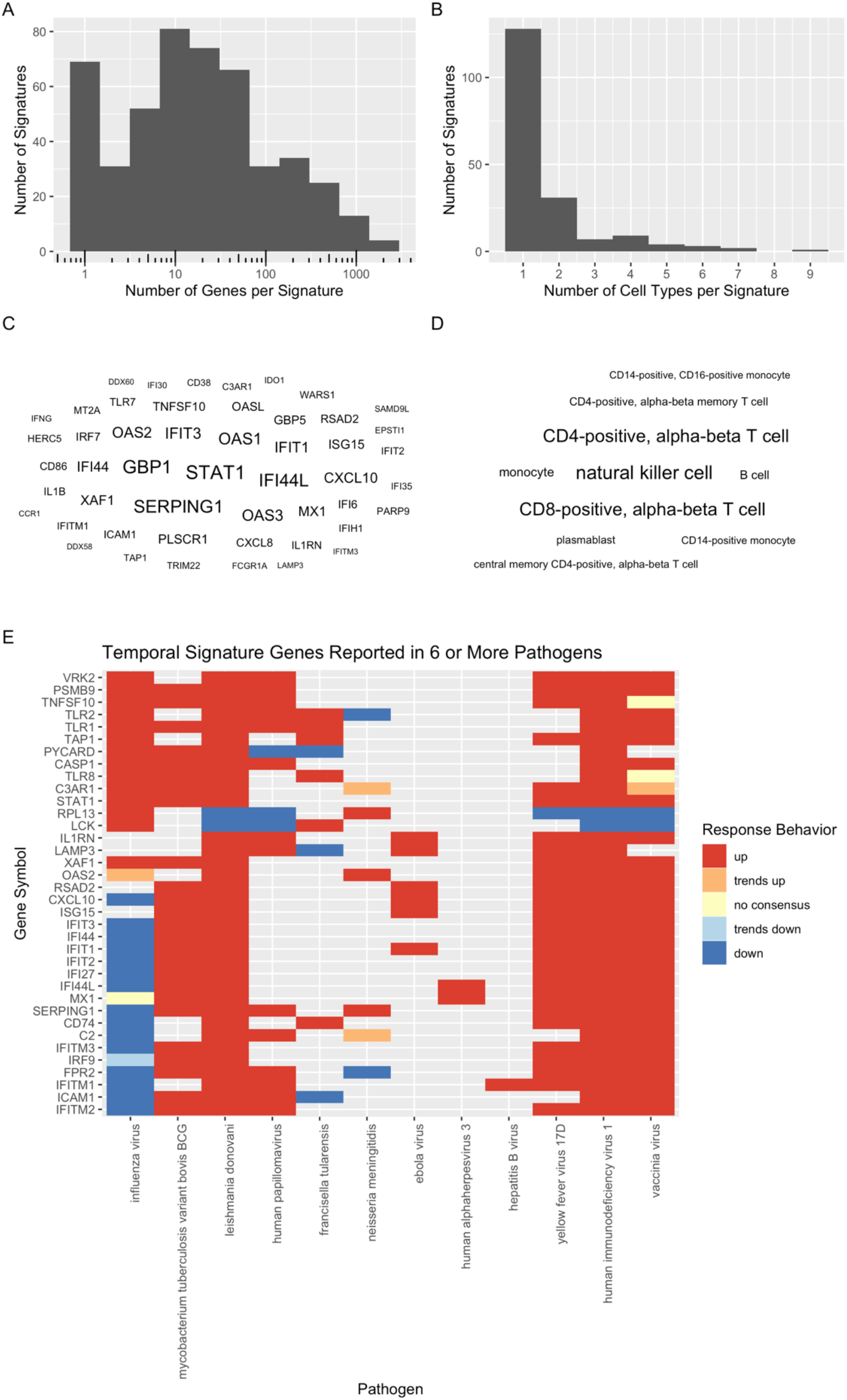
Summarization of HIPC Dashboard contents. Signature size distributions showing the number of response components across **A.** gene and **B.** cell type signatures. **C.** Word cloud of the top 50 most frequent gene symbols and **D.** top 10 most frequent cell types, where size corresponds to the total number of observations in the Dashboard. **E.** Heatmap of recurring genes across vaccines targeting different pathogens. Temporally associated genes in adult whole blood or PBMCs were filtered to those with signatures for six or more pathogens. Color indicates up (red) or down (blue) regulation. Genes with opposing directions in multiple studies were marked ‘trends up’ or ‘trends down’ according to the most common direction (or marked ‘no consensus’ for perfect ties).

The frequency with which cell types and genes are reported in the Dashboard offers insights about key players of the human immune response to vaccination. The most commonly reported genes are STAT1, a key mediator of immune response activated by cytokines and interferons; GBP1, an interferon induced gene involved in innate immunity; IFI44L, a paralog of Interferon Stimulated Protein 44, and SERPING1, a complement cascade protein **(Figure 3C)**. The most common cell types across pathogens and comparisons are NK cells, CD4+ T cells, and CD8+ T cells. **(Figure 3D)**. We searched the HIPC Dashboard data for genes with vaccination signatures across six or more pathogens and found a set of 36 associated genes across 12 target pathogens **(Figure 3E)**. Many are interferon stimulated genes, Toll-like receptors, or members of the complement cascade, potentially reflecting a common transcriptional program in response to many different vaccinations.

By design, our current implementation captures vaccination signatures as they are reported in the literature, but it does not include related methodological or statistical information regarding signature discovery (e.g. p-value cut-off) or provide analytical tools that can be applied to the curated signatures. Test statistics are not usually comparable across study designs, and we believe this information may give users a false sense that some signatures are more statistically reliable than others. We instead defer to the judgement of each study’s authors and their peer reviewers, and capture signatures as they were reported in each publication. We also caution that bias regarding the number of times particular genes or cell types were investigated might skew relationships in the Dashboard, thus precluding certain types of analyses. As a result, high level analytical tools have not been integrated into the Dashboard, although all of the signatures with full metadata can be downloaded to enable the use of third-party tools. Despite this, we believe the signatures available in the HIPC Dashboard will allow the research community to quickly query the literature and provide valuable comparisons and context for their own experimental results.

The number of genes and cell types captured reflects publications curated through January 2021, but we anticipate the HIPC Dashboard will undergo regular updates to accommodate new findings and additional domains of interest. The current implementation includes gene and cell type response components, as these represent the most commonly published signature types, but it will be valuable to also curate other response components, such as pathways, proteins, and metabolites. Additionally, we recognize that researchers may wish to compare a vaccine response against a particular pathogen to its corresponding disease response; it is easy to see how future iterations could expand the existing vaccine signature framework to capture signatures of infection. Based on our experience, we expanded the data model to include figure numbers or supplementary file annotations within publications as this can greatly simplify quality control during manual curation. We have provided links in the Dashboard to original sources wherever possible. We are also keen on exploring advancements in text-mining and artificial intelligence (AI), to assess how they can assist in automating signature identification and coding. To that end, the immune signatures in the HIPC Dashboard can be used as a data source for training/testing such AI solutions in the future.

In summary, we present the HIPC Dashboard (hipc-dashboard.org) to provide the vaccine research community with easy access to hundreds of published human systems vaccinology signatures. This resource will allow researchers to rapidly compare their own experimental results against existing findings that may otherwise be difficult to locate in the literature. This resource encourages the re-use of published results for advancing our understanding of human vaccine responses and provides a framework that can be extended to capture signatures from other types of immune exposures.

## Methods

### Manual Curation

The initial list of publications to curate into the HIPC Dashboard were derived from a PubMed search of papers matching the terms “Vaccine [AND] Signatures” or “Vaccine [AND] Gene expression”. Publications were further filtered to meet a set of inclusion criteria: (i) study involved human subjects, (ii) provided a comparison of a measurable change or correlation before and after vaccination (or challenge), and (iii) were reported as statistically significant. Signatures were excluded if they were missing directionality, or if they were derived from datasets external to the publication, to avoid redundancy. Two data curators manually collected a standard set of information from each study according to the designed data model (see **Figure 1**) and recorded it into a spreadsheet. Each signature was entered by one curator, and subsequently double-checked by the second curator. Table 3 shows a representative portion of the standardized information captured for each signature (12 data fields out of a total of 25).

Assays in the curated publications included gene expression analysis and measured changes in cell-type frequency and cell activation state. Each publication could give rise to any number and type of individual signatures. The signature content is centered on a list of biological response components (genes or cell types) that had a statistically significant change in the assay. These were designated as “response components” to capture different types of entities in a single standardized template column. For example, for gene expression assays, this was often a list of differentially expressed genes.

For genes, an initial manual curation process was applied to make a first pass at symbol standardization and detect any mistakes in copying. Gene names, symbols and or IDs (which may include HGNC symbols, Entrez IDs, Ensembl IDs etc.) were searched in turn against Panther, (www.pantherdb.org)^22^, using the “Functional classification viewed in gene list” search, followed if needed by searches against UniProt (www.uniprot.org)^23^ and NCBI (www.ncbi.nlm.nih.gov/search) until either a match or updated symbol was found; in the case of no match, the original representation was left unchanged. Any IDs that matched entries that were deprecated or defined as pseudogenes were removed from the curation.

### Data Standardization

The manually curated data required further steps to match terms encountered to their appropriate ontology representations. A number of the translations described below were orchestrated using an R script, which generated files ready for loading into the HIPC Dashboard.

#### Gene symbols

Outdated gene symbols and known aliases were translated to their current NCBI representation, which is the HGNC symbol in all but one case. The first pass of conversion used the function alias2SymbolUsingNCBI() from the Bioconductor limma package^24^ with the most recent available gene annotation file. This function returns either an exactly matching official symbol, or if none, the alias with the lowest EntrezID. We followed this by a second R package, HGNChelper^25^, which was able to resolve additional unmatched gene symbols to valid NCBI symbols. Genbank accession numbers were converted to gene symbols where possible using the org.Hs.egACCNUM2EG translation table which is part of the Bioconductor org.Hs.eg.db package^26^. Selected symbols still not matching NCBI names were investigated and corrected manually where possible after checking the original publications for context or for errors in transcription. Symbols for which no valid NCBI gene symbol was found, e.g. some pseudogenes, antisense, or uncharacterized genes, are not included in the HIPC Dashboard proper, since a requirement of the Dashboard framework is that all gene symbols must appear in the controlled vocabulary (NCBI/HGNC). However, these symbols are included in downloadable complete gene lists for each signature.

#### Cell types

Cell types as response components were first curated from the publications as published using a combination of cell type terms and additional descriptive terms, such as protein marker expression. This information was then mapped to a combination of Cell Ontology and Protein Ontology terms, according to a published model^16^. Note that cell types can appear in two different contexts, either as response components themselves, or as the cell type isolated for gene expression experiments. In some cases, additional information was provided which could not be mapped to an ontology term^16^. This type of information related to a wide variety of cell identification techniques and included the use of additional stains such as viability dyes or tetramer staining. For each set of terms, an entry was created in a lookup table by assigning (1) a parent cell type from the cell ontology, (2) mapping additional protein marker terms to the protein ontology, and then (3) separately retaining as free text descriptors not mapped to an ontology entry, such as tetramer specificity. Thus, the original entry was mapped to up to three descriptor columns, which can be combined as needed for display purposes. For a full, translated cell type, the displayed format is the cell ontology name followed by, if there are additional terms, the “&” symbol, followed by any PRO terms and then any free text.

#### Vaccines

Vaccine names collected from the literature were manually mapped to the most specific vaccine ontology term available. If a specific vaccine could not be found in VO, new terms were requested. Some examples of terms we included were VO_0004899 (2012-2013 seasonal trivalent inactivated influenza vaccine), VO_0003961 (ChAd63-KH vaccine, *Leishmania donovani*), VO_0004890 (gH1-Qbeta vaccine, novel pandemic-influenza), and VO_0004891 (CN54gp140□+□GLA, HIV-1).

#### Pathogens

The viral, bacterial or protozoan pathogens targeted by each vaccine are represented with terms from the NCBI Taxonomy. For the case of influenza vaccines, a table was created mapping vaccines by year of administration and type (e.g. trivalent or quadrivalent) to their seasonal viral components, unless otherwise indicated in the publication (e.g. for monovalent or specialized vaccines such as *Pandemrix*). For the few cases where the exact viral strain was not present in the NCBI Taxonomy, the closest more general term in the hierarchy was used. This mapping table was used to substitute in the actual viral pathogen names.

## Data Availability

Curated signatures are available on the HIPC Dashboard website (http://hipc-dashboard.org) and Github (see Code Availability for details). Files listing all response components for a signature can be downloaded from within individual observations in the Dashboard. The complete set of signature data can be downloaded from the GitHub repository at https://github.com/floratos-lab/hipc-dashboard-pipeline. This repository contains copies of (1) the original curated data sheets, (2) the response components in individual files, one per signature, (3) the response components in the Broad GMT format^8^, and (4) the actual tab-delimited Dashboard load files, in which the complete signature data is fully denormalized into an easy-to-parse format. Further details about the file formats are available on the GitHub project page. R session information for the Dashboard signature pre-processing pipeline is available on GitHub.

### Code Availability

Source curated data and mapping files (cell types, vaccine components, the NCBI gene file used, etc.), as well as the R code for the processing pipeline used to create the Dashboard submission files, are available on GitHub at https://github.com/floratos-lab/hipc-dashboard-pipeline. The data in this paper corresponds to pipeline and data version 1.2.1 in the pipeline GitHub repository. Code for the HIPC Dashboard web interface is available on GitHub at https://github.com/floratos-lab/hipc-signature.

### Supported web browsers

The HIPC Dashboard has been tested on recent versions of

- Chrome (Version 93.0.4577.63 (Official Build) (64-bit))
- Firefox (Version 92.0 (64-bit))
- Edge (Version 93.0.961.38 (Official build) (64-bit))

## Acknowledgements

This research was performed as a project of the Human Immunology Project Consortium (HIPC) and supported by the National Institute of Allergy and Infectious Diseases. This work was supported in part by NIH grants U19AI128949, U19AI118608, U19AI118626, and U19AI089992. This work was supported in part by the Canadian Institutes of Health Research [funding reference number FDN-154287]

## Author contributions

Conceptualization: AF, SHK; Methodology: KCS, DC, BKD, ZJ, RV, BP, REWH, AF, SHK; Software: KCS, ZJ, AF; Investigation: KCS, DC, BKD, EvdL, JW, ET, AA; Writing - Original Draft: KCS, DC, BKD, AF, SHK; Writing - Review and Editing: all authors

